# Fertilisation of agricultural soils with municipal biosolids: Part 1- Glyphosate and aminomethylphosphonic acid inputs to Québec field crop soils

**DOI:** 10.1101/2023.12.14.571753

**Authors:** Ariane Charbonneau, Marc Lucotte, Matthieu Moingt, J. C. Andrew Blakney, Simon Morvan, Marie Bipfubusa, Frédéric E. Pitre

## Abstract

Municipal biosolids (MBS) are suggested to be abundant, sustainable, inexpensive fertilisers, rich in phosphorus and nitrogen. However, MBS can also contain glyphosate and phosphonates that can degrade to AMPA. Glyphosate-based herbicides (HBG) are used in field crops all over the world. Most glyphosate generally degrades within a few weeks, mainly as aminomethylphosphonic acid (AMPA). AMPA is more persistent than glyphosate, and can accumulate from one crop year to the next. AMPA is phytotoxic even to glyphosate-resistant crops. The aims of this study were to assess whether applications of MBS significantly contribute to: 1) glyphosate and AMPA contents of agricultural soils, 2) trace metal elements contents, and 3) partial replacement of mineral fertilisation while maintaining similar yields. To this end, four experimental agricultural sites were selected in Québec (Canada). Soil samples (0-20 cm) were collected to estimate the as yet unmeasured contribution of BSM application to glyphosate and AMPA inputs in agricultural soils. MBS applied in 2021 and 2022 had mean concentrations of 0.69 ± 0.53 µg glyphosate/dry g and 6.26 ± 1.93 µg AMPA/dry g. Despite the presence of glyphosate and AMPA in MBS, monitoring of these two compounds in corn and soybean crops over two years showed no significant difference between plots treated with and without MBS applications. For the same site, yields measured at harvest were similar between treatments. MBS application could thus represent a partial alternative to mineral fertilisers for field crops, while limiting the economic and environmental costs associated with their incineration and landfilling. It is also an economic advantage for agricultural producers given the possibility of using fewer mineral fertilisers and therefore reducing the environmental impact of their use.

**Highlights:**

Municipal biosolids (MBS) offer significant potential for fertilising field crops
MBS contain significant amounts of both glyphosate and AMPA
Soil chemistry and crop yields were followed in MBS treated and untreated plots
Glyphosate and AMPA contents in soil do not increase with MBS treatment
MBS are a good substitute for mineral fertilisers leading to similarly crop yields

**Graphical abstract:** 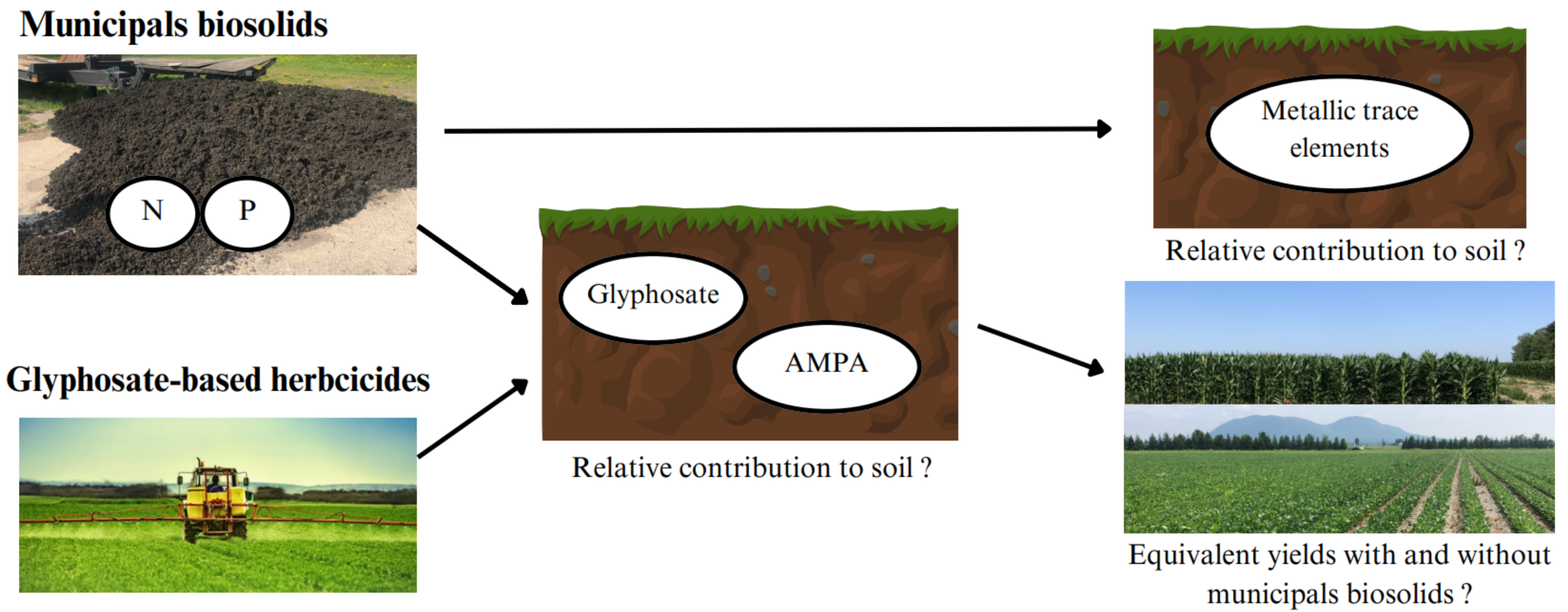

## 1.0 Introduction

The application of municipal biosolids (MBS) is now proposed in Québec’s waste management policy as an abundant and inexpensive source of phosphorous- and nitrogen-rich fertilisers for soils (Hébert, 2015). MBS comes from wastewater processing in treatment plants. Wastewater treatment not only represents high costs for communities but is also a major source of greenhouse gas (GHG). Methane (CH4) and nitrous oxide (N2O) are produced during wastewater treatment but also during disposal of the resulting waste, MBS (Brown *et al*., 2010). In fact, landfilling and incinerating MBS emits more GHG than land application (Brown *et al*., 2010). For this reason, the Québec Reference Centre for Agriculture and Agri-food (CRAAQ) recommends spreading MBS as residual fertiliser for field crops (Parent & Gagné, 2010). A number of studies claim that applying MBS provides yields as satisfactory as applying synthetic fertilisers (Vasseur *et al.,* 1999; Warman & Termeer, 2005). Although the application of MBS remains controversial, this practice has become increasingly popular with field crop producers in recent years in Québec and elsewhere (Hébert, 2015). In 2021, a total of 794 000 tonnes of MBS were generated in Québec, of which 346 000 tonnes (i.e. 44%) were spread in agricultural fields (Recyc-Québec, 2023).

Glyphosate-based herbicides (GBHs), intensively applied for over 40 years, are the most widely used in the world, with a production of 825 000 tons of active ingredient in 2014 (Benbrook, 2016). GBHs are used for weed control in agricultural settings, in conjunction with crops genetically modified to tolerate glyphosate, as well as in non-agricultural settings (Benbrook, 2016). In Québec, glyphosate represents 48% of agricultural sales with nearly 1700 tonnes (MELCCFP, 2023). Concerns about the risks to human and environmental health from the massive use of this class of herbicide continue to grow (Van Bruggen *et al*., 2018; Gillezeau *et al*., 2019). In 2015, the World Health Organization classified glyphosate as potentially carcinogenic to humans exposed to environmental doses (Van Bruggen *et al*., 2018). In contrast, the U.S. Environmental Protection Agency has found no risk to human health from glyphosate exposure at environmental doses (U.S. EPA, 2022). Dissipation of glyphosate in agricultural soils is generally rapid, on the order of a few weeks in Québec or under similar climatic conditions, but results, in parts, in the production of AMPA (Maccario, 2022; Silva *et al*., 2018; Silva *et al*., 2019; Struger *et al*., 2015). As it is the case around the world, glyphosate and aminomethylphosphonic acid (AMPA) have been detected in all sampled surface waters in Québec for over a decade (Giroux, 2019 & 2022). Field half-lives range from 5.7 to 40.9 days and 283.6 to 633.1 days for glyphosate and AMPA, respectively (Carretta *et al*., 2022; Lewis *et al*., 2016). AMPA may thus accumulate from one crop year to the next (Maccario, 2022). AMPA is also phytotoxic, even for plants genetically modified to tolerate glyphosate (Gomes *et al*., 2014; Smedbol *et al*., 2019). However, AMPA impacts on the environmental health have yet been sparsely studied (Van Bruggen *et al*., 2018).

In addition to the breakdown of glyphosate, a second important source of AMPA in the environment comes from the degradation of phosphonates (Botta *et al*., 2009; Grandcoin *et al*., 2017; Struger *et al*., 2015). These chelating agents are found in everyday household laundry and cleaning products (Jaworksa *et al*., 2002; May *et al*., 1986), flame retardants, anti-corrosion and anti-scaling products, and as complexing agents in the textile industry (Studnik *et al*., 2015; Nowack, 2003). Thus, the massive use of GBHs and phosphonates, including in urban areas, may explain glyphosate and AMPA omnipresence in the environment, particularly in surface waters (Giroux, 2019 & 2022), and wastewater treatment plants. (Botta *et al*., 2009). For example, in southern Ontario, Canada, using a municipal wastewater marker (the sweetener acesulfame), AMPA concentrations in streams were primarily linked to the use of GBH, but not to phosphonates in wastewater (Struger *et al*., 2015). In France, high concentrations of glyphosate in surface waters in urban areas after heavy rainfall have been attributed to the leaching of GBH residues from roadside and railroad tracks (Botta *et al*., 2009). In the same study, systematically high concentrations of AMPA in sewage water were reported and attributed to the partial degradation of phosphonates contained in detergents. However, glyphosate and AMPA contents are not routinely monitored in MBS production, or field application.

For MBS to be valued in the field, they must meet environmental quality criteria, particularly regarding chemical contaminants, pathogens, odors and foreign bodies (C-P-O-E) (Hébert, 2015). However, some contaminants, such as trace metals, are of great concern. Several studies have been carried out on the risks of trace metal contamination associated with the use of MBS on agricultural fields (Marcela *et al*., 2020; Perron & Hébert, 2008; Pepper *et al*., 2008). These generally show an increase in the contents of some trace elements, without however exceeding the regulatory limits of the countries where studies have been run. Furthermore, to our knowledge, only one study reports AMPA contents in MBS, namely Ghanem *et al*. (2007) in France, with values ranging from 1 to 30 µg of AMPA·g^-1^.

We tested the following hypotheses: 1) applying MBS can significantly contribute to glyphosate and AMPA contents in field crop soils, in addition to the ones resulting from GBHs applications, 2) applying MBS does not increase the risk of soil contamination by trace metal elements, and 3) yields will be equivalent between plots treated with and without MBS. In the present study, total fertiliser inputs between plots were equal. Plots with MBS were fertilized using one MBS application and a complementary application of mineral fertilisers to match the mineral fertilization of plots without MBS application.

## 2.0 Materials and Methods

### 2.1 Study sites & experimental design

The experiment was established in eight open fields at four agricultural sites in Quebec, Canada (Fig. 1). Sites were selected based on similar soil types (silty-clay), use of corn-soybean rotations, use of glyphosate-based herbicides, as well as agricultural practices including conventional production methods and direct seeding. At each site in 2021 and 2022, two fields were prepared with a randomized complete block design with four replications, where each block was randomly divided into MBS treated and untreated plots (Fig. 2). The numbers in the figure correspond to the plot numbers. In 2021, one field was seeded with Roundup Ready soybean, while the second was seeded with Roundup Ready corn. In 2022, following the same experimental design, crops were alternated, establishing soybean-corn and corn-soybean rotations over 2021 and 2022 at each site. Therefore, in total we had 384 samples: 2 years * 4 sites * 2 crop soils * 2 treatments * 4 blocks * 3 sampling campaigns.

**Figure 1.**
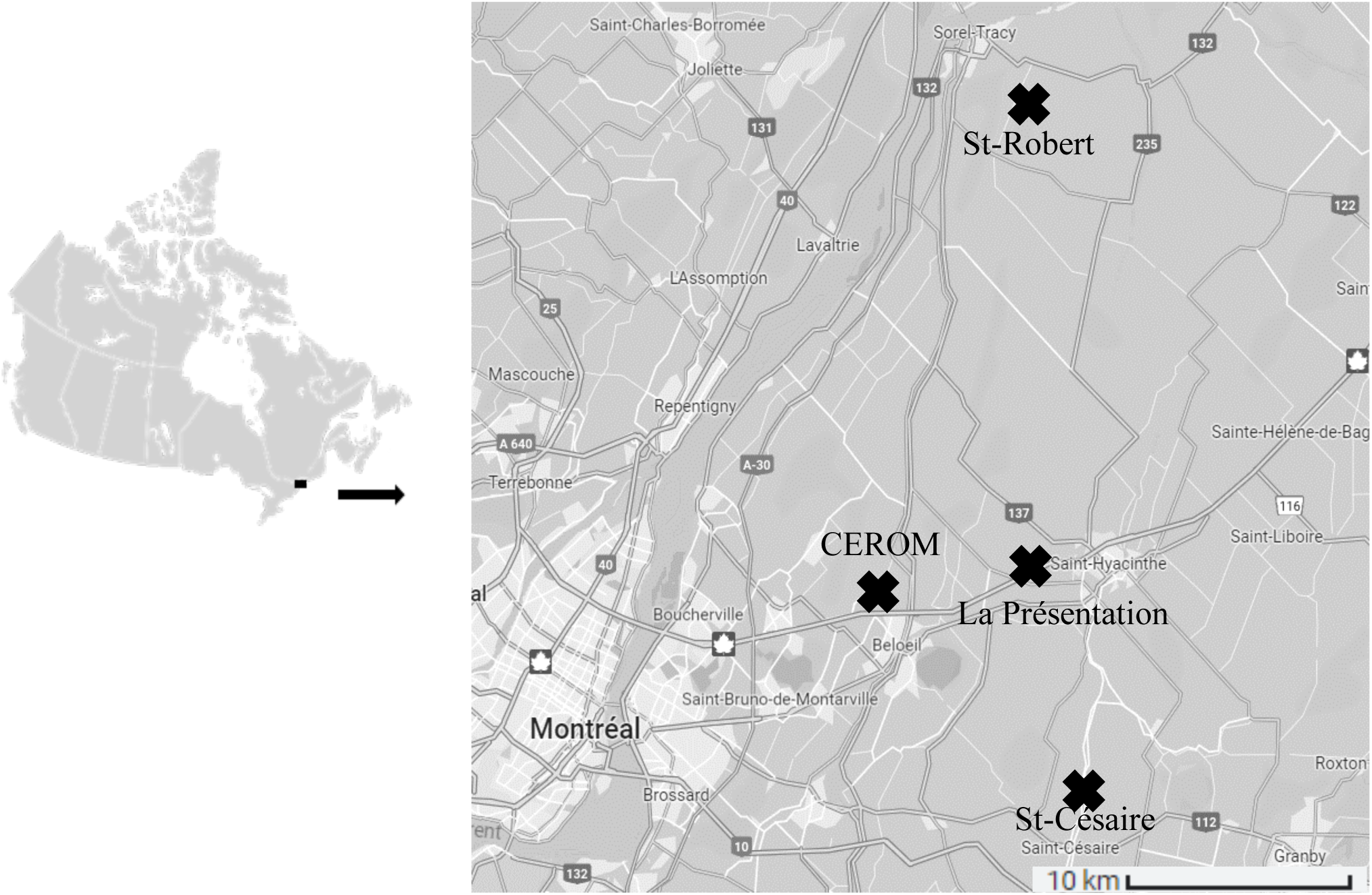
Field sites were located in East Montérégie, Québec; Saint-Robert (46°00ߴ N, -72°96ߴ W), La Présentation (45°62ߴ N, -73.05ߴ W), Saint-Césaire (45°41ߴ N, -72°96ߴ W), and Saint-Mathieu-de-Beloeil (45°58ߴ N, -73°24ߴ W)

**Figure 2.**
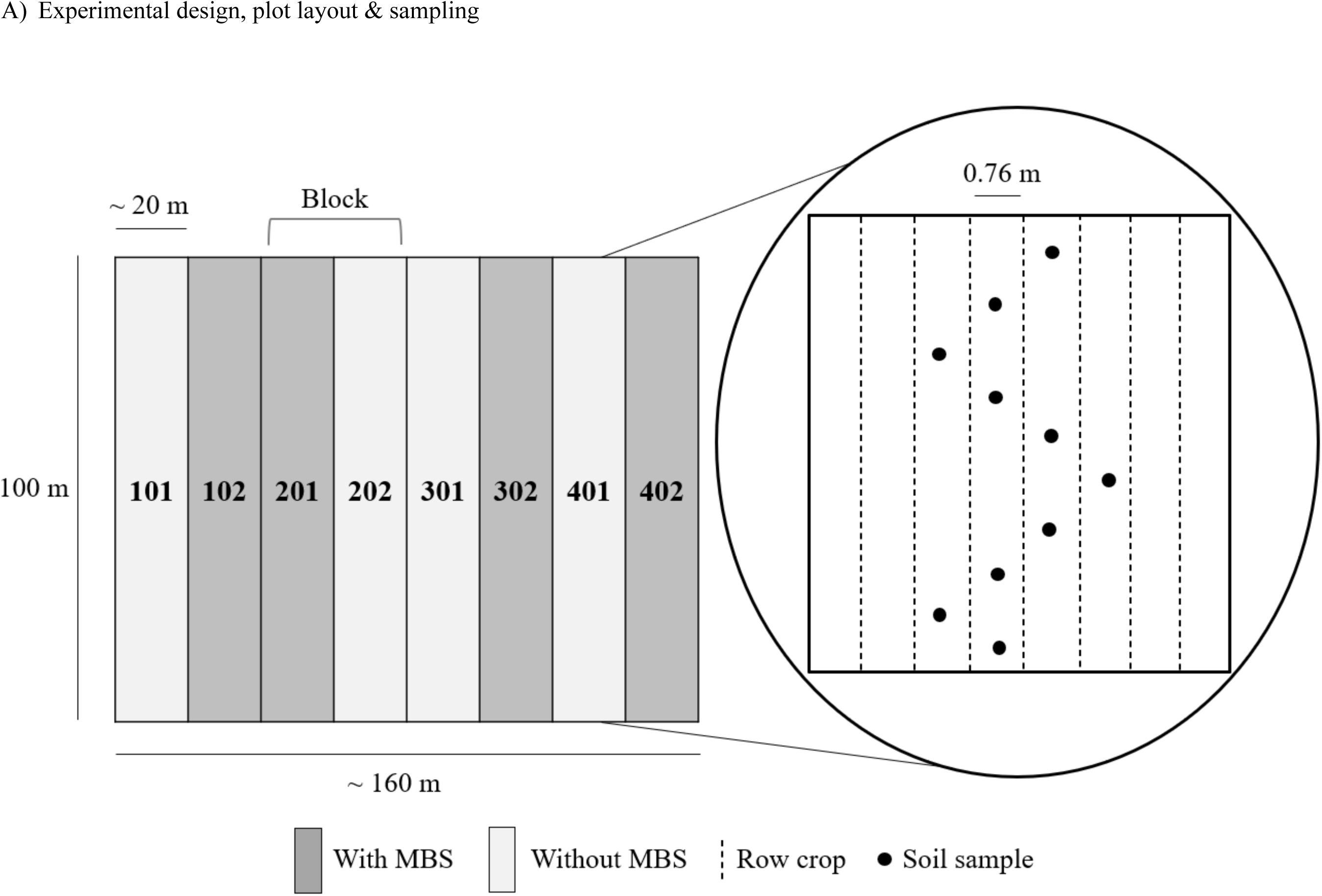
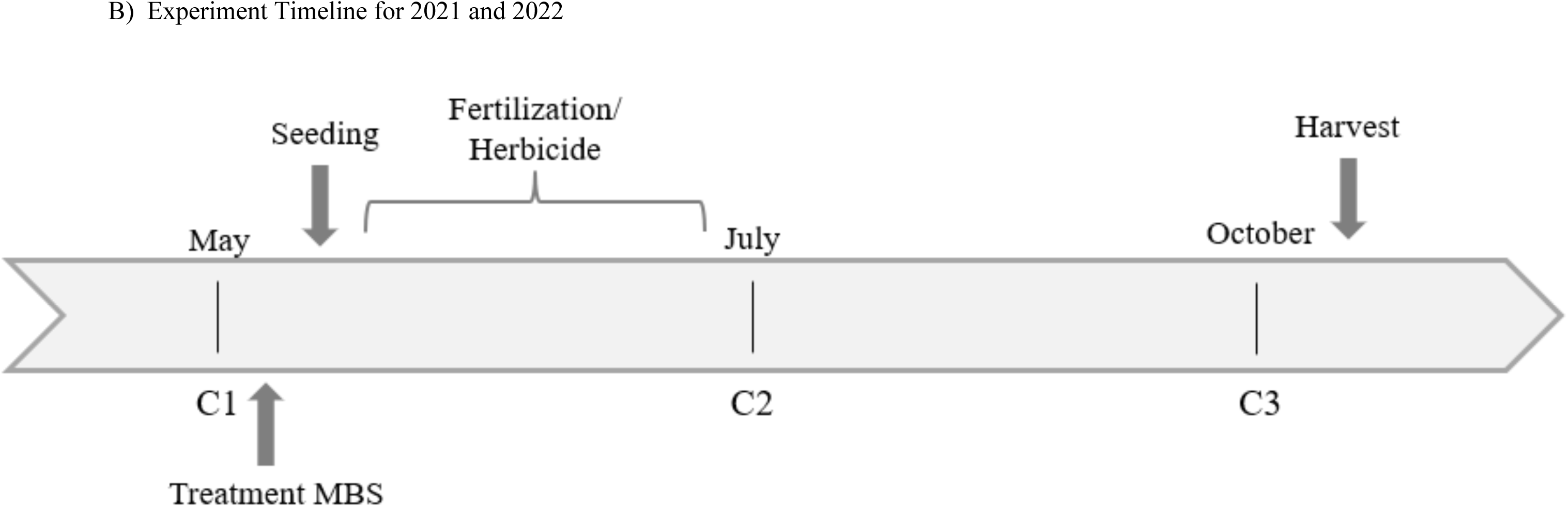
A) Experimental design of the plots set up on 4 study sites, plot layout & sampling, B) Experiment Timeline for each site.

### 2.2 Crop management and sampling

Crops at all sites were grown and maintained following the recommendations of the CRAAQ standard agricultural practices. Corn and soybean were seeded from May 7^th^ to May 14^th^, 2021 and 2022, except at CÉROM where soybean was seeded June 14^th^ in 2022, due to weather condition (Fig. 2B). MBS was obtained from a regional wastewater treatment plant and applied to the experimental plots in early May before seeding (Fig. 2B). MBS applied to the field were from the C2-P2-02-E2 category. The different MBS categories come into play regarding the type of culture, and the separation distances between application site and neighbors. The doses of MBS were determined for each site in accordance with the local environmental legislation (i.e. the “*Règlement sur les exploitations agricoles”)*, which stipulates that Québec farms are required to respect a maximum phosphorous dosage as determined by the Ministry of the Environment (Parent & Gagné, 2010). Therefore, MBS dosage was calculated based on, first, the estimated phosphorous contents of the MBS deriving from the average phosphorus contents measured in the MBS of the wastewater treatment plant over the years 2019 to 2021, and second, the phosphorus requirements of each crop according to the CRAAQ guidelines (Parent & Gagné, 2010). Fertiliser nutrient inputs (N-P-K) were equivalent between the plots with and without MBS application. However, control plots were solely fertilized using mineral fertilisers whereas treated plots were mainly fertilized with MBS and complementary mineral fertilisers to fulfill the remaining nutritional needs. Annual average temperature (°C) and total precipitation (mm) per day, for each site are reported in Table 1.

**Table 1.** Environmental variables of the four experimental sites. For each site, soil texture was determined at the *Institut de Recherche et de Développement en Agroenvironnement* (IRDA) with soil samples from 0-20 cm depth taken in the Fall of 2020; average precipitation, average temperature and 30-year average from May to October were all taken from the nearest weather station to the site (MELCCFP, 2022).

| Site | Soil Texture | Precipitation (mm)<sup>a</sup> |  |  | Temperature (°C)<sup>a</sup> |  |  |
| --- | --- | --- | --- | --- | --- | --- | --- |
|  |  | 2021 | 2022 | 1991-2020 | 2021 | 2022 | 1991-2020 |
| CÉROM | Heavy clay<br>(Clay 72%, Silt 28%, Sand 0%) | 73.0 | 115.3 | 102.1 | 17.7 | 16.9 | 16.3 |
| La<br>Présentation | Clay loam<br>(Clay 30%, Silt 34%, Sand 36%) | 73.0 | 115.3 | 102.1 | 17.7 | 16.9 | 16.3 |
| Saint-Césaire | Silty clay loam<br>(Clay 35%, Silt 46%, Sand 19%) | 79.9 | 110.1 | 106.5 | 17.7 | 17.0 | 16.5 |
| Saint-Robert | Silty clay loam<br>(Clay 35%, Silt 50%, Sand 15%) | 81.5 | 108.4 | 99.9 | 18.1 | 17.5 | 16.7 |

<sup>a</sup>, Government of Québec, Ministry of the Environment, the Fight Against Climate Change, Wildlife and Parks, 2022.

In 2021 and 2022, each plot was sampled three times per year (Fig. 2C) Campaign 1 (C1), used as a baseline control, was sampled before MBS application, seeding and the initial use of glyphosate-based herbicide, whereas campaign 2 (C2), was performed during the growing season, 7 to 10 days following the last GBH application. Finally, the third campaign (C3) took place before harvesting. During each campaign, 10 soil cores per plot, ∼5 m apart, were randomly sampled across the plot using a manual corer at 0-20 cm depth (Fig. 2B). All 10 cores were pooled to form composite sample for each plot (Fig. 2B) to obtain representative values for soil chemistry in the given plot as suggested by (Khiari, 2014). Each sample core was geo-referenced for subsequent sampling campaigns over both 2021 and 2022. In the field, each sample was manually homogenized in plastic bags, subdivided in identified containers and placed in coolers containing ice-packs, before being stored at -20°C back at the university until processing. Soil samples were then freeze-dried, ground and sieved (2 mm pore size).

### 2.3 Carbon, nitrogen, glyphosate and AMPA contents

The percentage of organic carbon and total nitrogen of the soil samples by weights were measured using a CE-Instruments NC2500TM elemental analyzer, with a relative precision of ± 5 % (1σ), corrected for atomic weight. We estimated the percentage of organic matter in the soil for C3 samples, from 2021 and 2022, by multiplying the percent of organic carbon by a factor of 2.0. Although the factor of 1.72 is routinely used, this approach often underestimates the organic matter content of soils (Pribyl, 2010). A factor of 2.0 is found to generate results closer to reality in the majority of situations (Pribyl, 2010).

A slightly modified procedure from Maccario *et al*. (2022) and Samson-Brais *et al*. (2022) was used to analyse glyphosate and AMPA contents. Briefly, an extraction solution was prepared by mixing 34.5 mL of NH4OH (28-30%) with 13.6 g de KH2PO4 and adjusting the final volume to 2 L. A 40 mL volume of the extraction solution was transferred to a Falcon tube containing 5 g of soil sample. The Falcon was mixed on a vortex for 30 seconds before being placed on a rotation wheel at 200 rpm for 45 minutes. A Varian GC 3800 gas chromatograph fitted with a Zebron ZB-1 (30 m 0.25 mm ID, 0.25 m) was used to analyze the samples. The chromatograph conditions used to detect glyphosate and AMPA were as follows: 280 °C injector temperature; 300 °C detector temperature, and an oven temperature program, 70 °C (hold for 1 min, 1 °C ·min^-1^ at 84 °C, 4 °C ·min^-1^ at 120 °C, 80 °C·min^-1^ at 250 °C, hold for 5 min, for a total cycle time of 30.63 min).

To minimise uncertainty in chromatograph measurements, GC-ECD performance parameters were checked daily to ensure that they were suitable for glyphosate/AMPA analysis. The detection limit and the quantification limit were determined on the basis of the basis described by Mocak *et al*. (1997). The calculated detection limits and quantification limits were 0.02 µg·g^-1^ and 0.05 µg·g^-1^, and 0.03 µg·g^-1^ and 0.09 µg·g^-1^ for glyphosate and AMPA, respectively (Samson-Brais *et al*., 2022). Six point calibration curves showed good linearity for both analyses (r^2^ > 0,95; p < 0,0001 for glyphosate and AMPA) within the expected concentration range. With regard to sample quantification, each batch of samples included a standard curve made up of five standards (0, 0.1, 0.2, 0.3, 0.4 and 0.6 µg·g^-1^, and 0, 0.2, 0.4, 0.6, 0.8 and 1.2 µg·g^-1^ for glyphosate and AMPA, respectively) in the same matrix as the samples.

### 2.5 Elementary soil analysis

Elementary analyses were performed on soil samples from C1 and C3 for both 2021 and 2022. The analyses were carried out by the agri-environmental analysis laboratory at the *Institut de Recherche et de Développement en Agroenvironnement* (IRDA). This laboratory is accredited by the MELCCFP and by the Ministry of Agriculture, Fisheries, and Food of Québec according to the international standard ISO/CEI 17025. The Mehlich Ⅲ method was used to extract all soil elements. Detection limits were 3 mg·kg^-1^ for phosphorus (P), potassium (K) and aluminium (Al), 5 µg·g^-1^ for calcium (Ca), 2 µg·g^-1^ for magnesium (Mg) and iron (Fe), 0.06 µg·g^-1^ for boron (B), molybdenum (Mo), nickel (Ni), cobalt (Co) and lead (Pb), 0.1 µg·g^-1^ for copper (Cu) and zinc (Zn), 0.5 µg·g^-1^ for manganese (Mn) and sodium (Na), and 0.02 µg·g^-1^ for cadmium (Cd) and chrome (Cr). The quantification limits were 10 µg·g^-1^ for P, K and Al, 15 µg·g^-1^ for Ca, 5 µg·g^-1^ for Mg and Fe, 0.2 µg·g^-1^ for B, Mo, Ni, Co and Pb, 0.4 µg·g^-1^ for Cu and Zn, 2 µg·g^-1^ for Mn and Na, and 0.05 µg·g^-1^ for Cd and Cr.

### 2.6 Yield measurements

Yield and grain moisture measurement were taken at harvest, directly in the field where each producer was equipped with a yield sensor. During the two years of the study, yields and grain moisture were measured once each year at the 4 sites at crop harvest. Yields were then adjusted to 15% moisture to enable comparison between control and MBS-treated plots. Yields are expressed in tonnes per hectare and moisture content is expressed as a percentage (%).

### 2.7 Statistical analyses

In order to use samples in our analyses where the glyphosate or AMPA contents were below the detection limit, or between the detection limit and the quantification limit, we assigned arbitrary values. Specifically, glyphosate or AMPA contents below the detection limit were assigned a (detection limit)/2 (i.e. 0.01 µg·g^-1^and 0.02 µg·g^-1^ for glyphosate and AMPA, respectively), while samples with contents between the detection limit and quantification limit were assigned a (quantification limit)/2 value (i.e. 0.03 µg·g^-1^ and 0.05 µg·g^-1^ for glyphosate and AMPA, respectively).

All statistical analyses were performed on R (4.2.1. R Core Team, 2022) under a Windows 10 operating system. For statistical comparisons, the significance level was set a *p value* < 0.05. The *lme4* package (Bates *et al*., 2015) was used to run all the following models, while the marginal r-squared values for the mixed models were calculated based on the method of Nakagawa *et al*. (2017). We fitted a generalized linear mixed model to predict glyphosate and AMPA as a function of the interactions between treatment, sampling campaign, and year, for each crop type. The model used a gamma distribution with a logarithmic link function, which is appropriate when the response variance increases with the average, and the response variable is positive, close to zero, and continuous. Next, we fitted a linear mixed model to predict organic matter and trace metal contents as a function of the interaction between treatment and sampling campaign for each crop type. Finally, we fitted a random mixed model to predict crop yields as a function of the interaction between treatment and sampling year for each crop type. All models include the effect of site and block as random effects. The *ggplot2* package (Wickham *et al*., 2016) was used to generate the graphs presented in the article.

## 3.0 Results

### 3.1 Chemical characteristics of MBS

Chemical characteristics of the applied MBS were provided by the waste management and reclamation company that delivered the product. Average N, P, and K, for the MBS from 2021 and 2022 were 12.86 ± 0.02, 15.56 ± 0.21 and 1.84 ± 0 kg·t^-1^, respectively. MBS were 66% organic matter (OM) (on a dry basis) in 2021 and 2022. Despite the high OM content, MBS application did not significantly impact OM contents in either MBS-treated soil corn plots (*p* > 0.05; R^2^ = 0.58) or soybean plots (*p* > 0.05; R^2^ = 0.49) in the 0-20 cm study area at the time of sampling campaign 3 in 2021 and 2022. After two years of MBS application, OM percentages remained similar in soils from plots with and without MBS.

### 3.2 Glyphosate and AMPA contents in MBS

Glyphosate and AMPA contents (Table 2) varied, sometimes considerably, between and within sites, highlighting the heterogeneity of MBS composition. This variation did not correspond to the delay between the date of delivery and sampling of the MBS at each site, as this delay was different from one site to another. No pattern could be observed between the delivery and sampling time and the glyphosate and AMPA contents measured in the MBS. MBS applied in 2021 and 2022 had mean concentrations of 0.69 ± 0.53 µg glyphosate·dry g^-1^ and 6.26 ± 1.93 µg AMPA·dry g^-1^. The maximum contents measured are 1.27 ± 0.60 µg glyphosate·dry g^-1^ and 8.48 ± 1.65 µg AMPA·dry g^-1^.

**Table 2.**
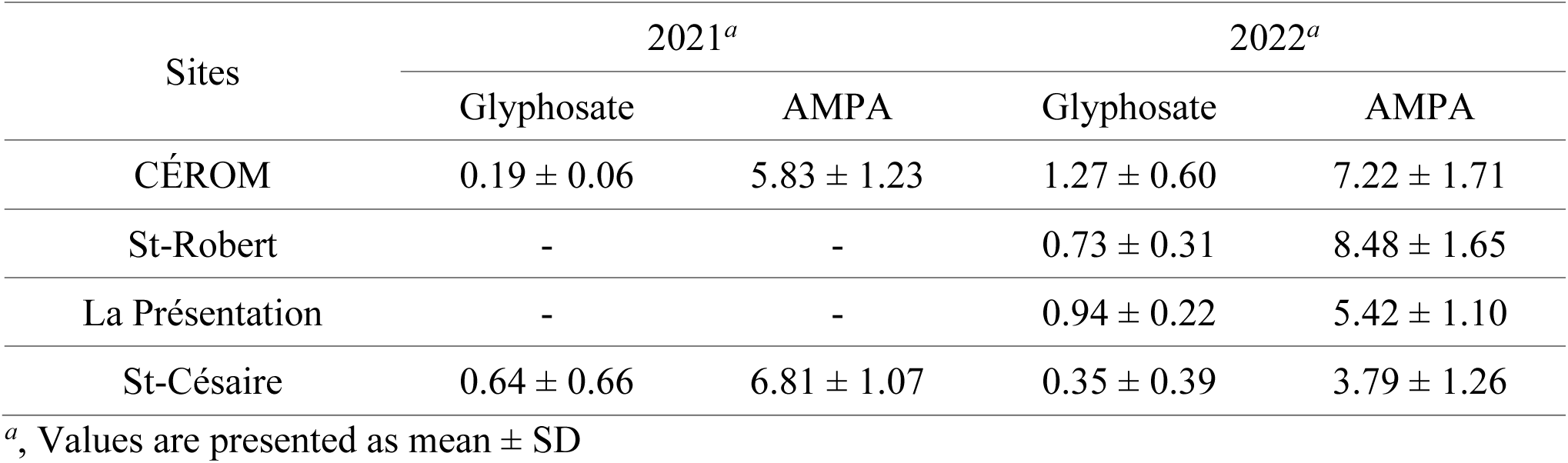
Glyphosate and AMPA contents in MBS in 2021 and 2022 at the four study sites. For each site: the mean and standard deviation of glyphosate and AMPA contents were determined (four subsamples per MBS pile).

| Sites | 2021<sup>a</sup> |  | 2022<sup>a</sup> |  |
| --- | --- | --- | --- | --- |
|  | Glyphosate | AMPA | Glyphosate | AMPA |
| CÉROM | 0.19 ± 0.06 | 5.83 ± 1.23 | 1.27 ± 0.60 | 7.22 ± 1.71 |
| St-Robert | - | - | 0.73 ± 0.31 | 8.48 ± 1.65 |
| La Présentation | - | - | 0.94 ± 0.22 | 5.42 ± 1.10 |
| St-Césaire | 0.64 ± 0.66 | 6.81 ± 1.07 | 0.35 ± 0.39 | 3.79 ± 1.26 |

<sup>a</sup>, Values are presented as mean ± SD

### 3.3 Glyphosate and AMPA contents in soils

Glyphosate contents 2021 and 2022 soil samples (n = 384) ranged from below the detection limit to 1.04 µg·g^-1^, while AMPA contents varied from below the detection limit to 1.69 µg·g^-1^. At least one of the two compounds was detected in 94% of the samples, and both glyphosate and AMPA were detected in 85% of the samples. At each sampling time (C1, C2, C3) in both years of the study, no significant difference was observed in the soil between plots of corn with and without MBS application at any of the four sites with regard to glyphosate contents (*p* > 0.05; R^2^ = 0.40, Fig. 3), or AMPA contents (*p* > 0.05; R^2^ = 0.33, Fig. 4). There were also no significant difference in glyphosate contents (*p* > 0.05; R^2^ = 0.52, Fig. 3), or AMPA contents (*p* > 0.05; R^2^ = 0.38, Fig. 4) measured in any of the soybean fields. This result is the same at each site taken separately (Fig. S1-S4). MBS application therefore does not appear to significantly increase glyphosate and/or AMPA contents in field crop soils. Rather, glyphosate and AMPA contents measured in the soils appeared mainly due to GBH applications.

**Figure 3.**
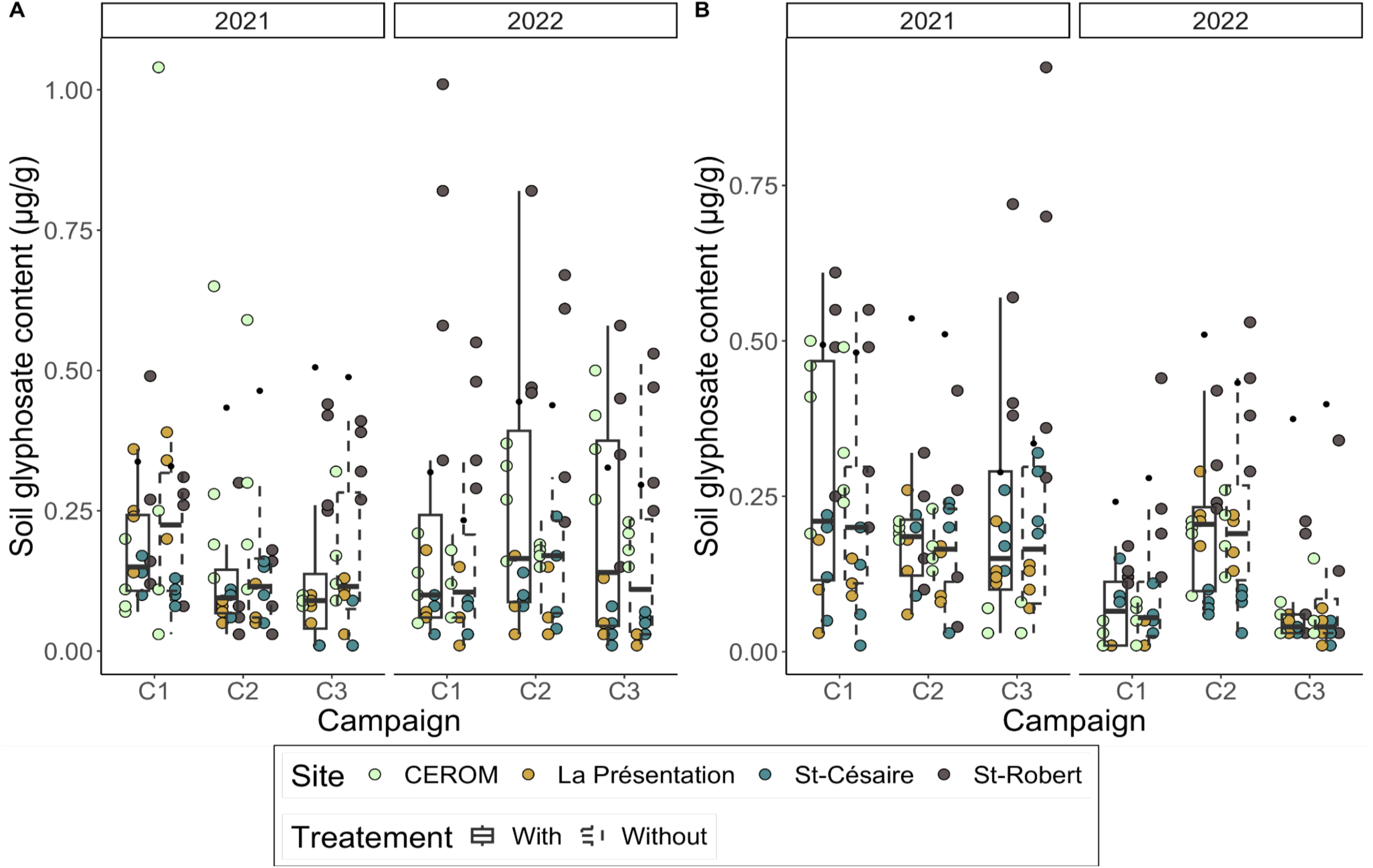
Soil glyphosate content (µg/g) in (A) corn and (B) soybean fields measured at each sampling campaign in 2021 and 2022. In each plot, 10 soil cores were sampled with a probe at a depth of 0-20 cm. The 10 cores were pooled to form composite samples for each plot, which is what each point in the figure corresponds to. A generalized linear mixed model predicted glyphosate as a function of the interactions of the treatment, sampling campaign and year for each crop type (*p* < 0.05). Boxplots represent median, and 25 and 75 % quantile. Dots represent glyphosate contents measure in the soil (n = 4 for a total of n = 384). Colours represents site, solid lined boxplots represent MBS treatment, and dotted lines represent plots without MBS treatment.

**Figure 4.**
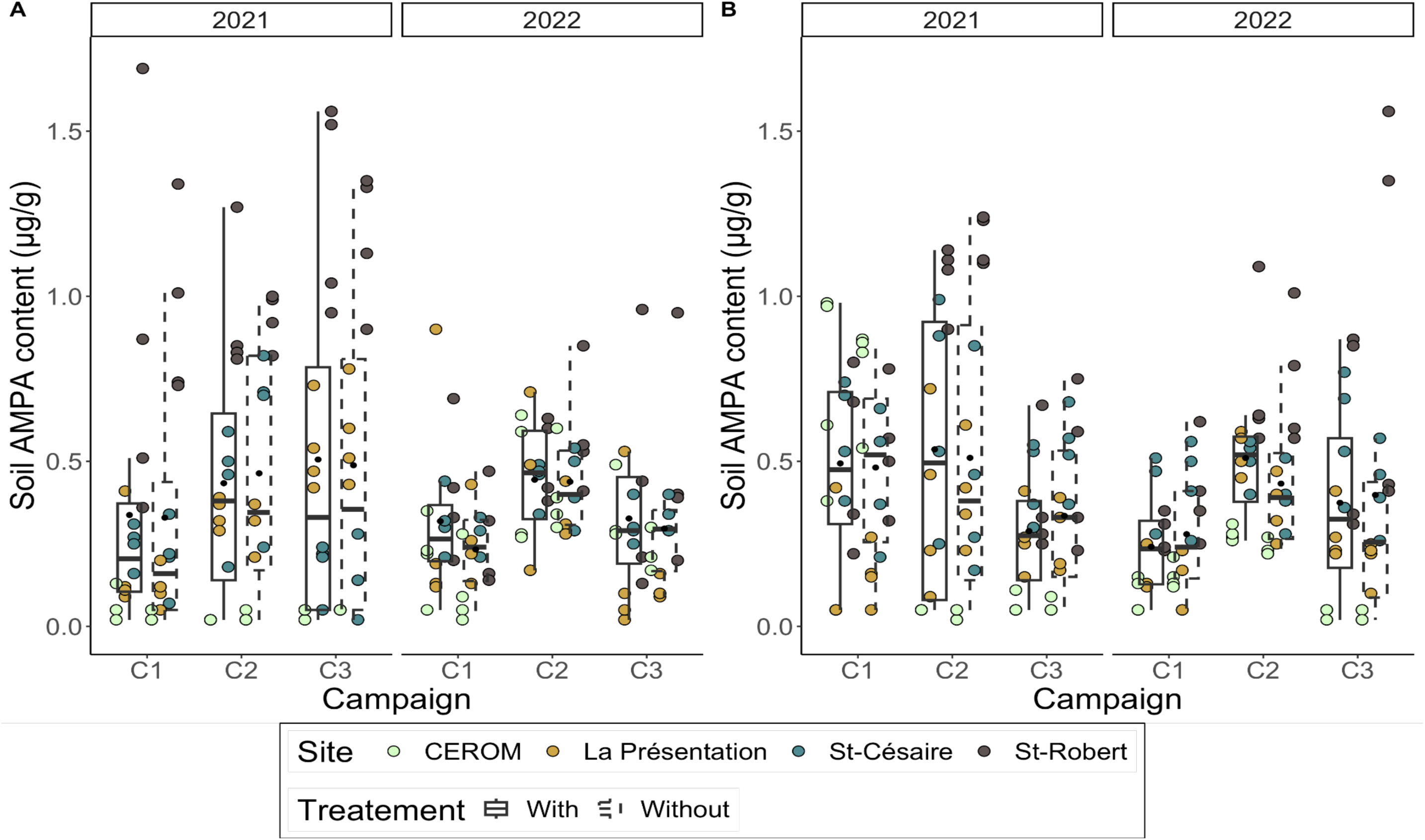
Soil AMPA content (µg/g) in (A) corn and (B) soybean fields measured at each sampling campaign in 2021 and 2022. In each plot, 10 soil cores were sampled with a probe at a depth of 0-20 cm. The 10 cores were pooled to form composite samples for each plot, which is what each point in the figure corresponds to. A generalized linear mixed model predicted glyphosate as a function of the interactions of the treatment, sampling campaign and year for each crop type (*p* < 0.05). Boxplots represent median, and 25 and 75 % quantile. Dots represent glyphosate contents measure in the soil (n = 4 for a total of n = 384). Colours represents site, solid lined boxplots represent MBS treatment, and dotted lines represent plots without MBS treatment.

### 3.4 Elementary contents of MBS

Elementary analyses of MBS show that this product may contain significant amounts of trace metals (Cu, Zn, Ni, Cd, Cr, Pb). But the application of MBS over two consecutive years had no impact on Cu contents (*p* > 0.05; R^2^ = 0.99), Zn (*p* > 0.05; R^2^ = 0.69), Ni (*p* > 0.05; R^2^ = 0.83), Cd (*p* > 0.05; R^2^ = 0.87), Cr (*p* > 0.05; R^2^ = 0.76) and Pb (*p* > 0.05; R^2^ = 0.93) in corn and soybean soils (Figure S5). No significant difference was observed in elementary analyses between plots with and without MBS at the four study sites. Furthermore, all elementary contents of MBS met the regulatory standards (Hébert, 2015).

### 3.5 Crop yields

No significant differences were observed in crop yields between plots with and without MBS application at any of the four sites for both corn crops (*p* > 0.05; R^2^ = 0.51) and soybean crops (*p* > 0.05; R^2^ = 0.46) in 2021 or 2022 (Table 3). Yields in plots treated with MBS were similar to those in control plots and were in line with typical yields in the study region. Yields in 2021 were higher than the regional average;10.10 t·ha^-1^ and 3.40 t·ha^-1^ for corn and soybean crops, respectively (ISQ, 2023), with the exception of corn yields in control plots at CÉROM site (9.96 t·ha^-1^) (Table 3). Yields for 2022 were also higher than the regional average, with 10.77 t·ha^-1^ and 3.39 t·ha^-1^ for corn and soybean, respectively (ISQ, 2023) (Table 3).

**Table 3.** Corn yields (A) and soybean yields (B) in plots with and without MBS application. Yields were measured at harvest in 2021 and 2022 for corn and soybean in tonnes/hectare. A linear mixed model predicted yield as a function of the interaction between treatment and year for each crop (*p* < 0.05).

| A | Sites | 2021<sup>a</sup> |  | 2022<sup>a</sup> |  |
| --- | --- | --- | --- | --- | --- |
|  |  | with MBS | without MBS | with MBS | without MBS |
|  | CÉROM | 10.25 ± 0.09 | 9.96 ± 0.38 | 12.61 ± 0.39 | 12.03 ± 0.43 |
|  | St-Robert | 10.52 ± 0.64 | 10.86 ± 0.46 | 11.61 ± 0.55 | 11.16 ± 0.66 |
|  | La Présentation | 12.76 ± 0.27 | 12.44 ± 0.57 | 13.04 ± 0.33 | 12.54 ± 0.40 |
|  | St-Césaire | 12.73 ± 0.39 | 12.89 ± 0.31 | 11.39 ± 0.10 | 11.66 ± 0.42 |
|  | Average yield of the region<sup>b</sup> | 10.10 |  | 10.77 |  |

| B | Sites | 2021<sup>a</sup> |  | 2022<sup>a</sup> |  |
| --- | --- | --- | --- | --- | --- |
|  |  | with MBS | without MBS | with MBS | without MBS |
|  | CÉROM | 4.26 ± 0.28 | 4.26 ± 0.25 | 3.60 ± 0.68 | 4.25 ± 0.56 |
|  | St-Robert | 3.84 ± 0.42 | 3.69 ± 0.47 | 3.93 ± 0.47 | 4.07 ± 0.43 |
|  | La Présentation | 4.46 ± 0.04 | 4.42 ± 0.22 | 5.54 ± 0.10 | 5.47 ± 0.15 |
|  | St-Césaire | 4.83 ± 0.15 | 4.57 ± 0.19 | 3.57 ± 0.28 | 3.69 ± 0.19 |
|  | Average yield of the region<sup>b</sup> | 3.50 |  | 3.30 |  |

<sup>a</sup>, Values are presented as mean ± SD
<sup>b</sup>, Institut de la statistique du Québec (ISQ)

## 4.0 Discussion

### 4.1 Glyphosate and AMPA contents were higher in MBS than in soils

One objective of this study was to assess whether MBS applications increased glyphosate and/or AMPA contents in soil. Several studies have quantified glyphosate and AMPA contents in water from wastewater treatment plants (Kolpin *et al*., 2006; Botta *et al*., 2009; Struger *et al*., 2015) and in different residual fertilisers (Ghanem *et al*., 2007), but never in agricultural soils following the application of MBS from wastewater treatment plants. Glyphosate contents in MBS (0.19 ± 0.06 µg·g^-1^ to 1.27 ± 0. 60 µg·g^-1^; Table 2) were generally higher to contents reported for agricultural soils in North America (0.07 ± 0.10 µg·g^-1^) (Maccario *et al*., 2022), South America (0.08 ± 0.09 µg·g^-1^ to 0.35 ± n.a. *^a^* µg·g^-1^) (Giard *et al*., 2022; Bento *et al*., 2019) and Europe (0.10 ± n.a. *^a^* µg·g^-1^) (Silva *et al*., 2018). Moreover, AMPA contents measured in MBS (3.79 ± 1.25 µg·g^-1^ to 8.48 ± 1.65 µg·g^-1^; Table 2) were well above those previously reported in the literature for agricultural soils (0.15 ± n.a. *^a^* µg·g^-1^ to 1.5 ± n.a. *^a^* µg·g^-1^) (Maccario *et al*., 2022; Giard *et al*., 2022; Silva *et al*., 2018 and Bento *et al*., 2019). To our knowledge, only one study in France has presented contents in MBS from different wastewater treatment plants ranging from 1 to 30 µg of AMPA·g^-1^ (Ghanem *et al*., 2007). Contents measured in MBS from the present study fall within this range (Table 2; Ghanem *et al*., 2007). Moreover, a study by Kolpin *et al*. (2006) conducted at 10 sites across the USA reported that over 65% of their sewage plant water samples contained AMPA, while over 15% of the samples contained glyphosate. The majority of these concentrations were below 1 µg·L^-1^ for both compounds, except for three samples for glyphosate and nine samples for AMPA out of the 40 samples (Kolpin *et al.,* 2006). Depending on the wastewater treatment process, it is possible to observe different glyphosate and AMPA contents in MSB produced by different wastewater treatment plants. Wastewater treatment plants in Quebec are either mechanized or “pond” type stations (Vigneux *et al.,* 2016). Our study was limited to measuring glyphosate and AMPA contents in MBS applied in the Québec agricultural region from a single local wastewater treatment plant where no drying is used. Additionally, as previously mentioned, all MBS belong to a specific quality class (i.e. C2-P2-02-E2) according to the four environmental criteria allowing their use as fertilisers in field crops.

Despite the quantifiable presence of glyphosate and high AMPA contents in MBS used in this study, both glyphosate and AMPA contents in soils from plots receiving MBS applications were not different from those measured in un-treated control plots (Fig. 3 & 4). This is mainly due to the relatively small quantities of MBS that could be applied (∼4 tonnes wet weight/hectare) in order to comply with the maximum phosphorus inputs allowed under local fertilizing regulations (Parent & Gagné, 2010; Hébert, 2015). Indeed, the four sites in our study were characterized with a high phosphorous saturation index (Table S1 et S2). Therefore, the amounts of glyphosate from an application of GBH at the rate of 1.67 L·ha^-1^ with 540 g·L^-1^ active ingredient are of the order of 900 g·ha^-1^. Knowing that the glyphosate molar mass is 169.97 g·mol^-1^, we obtain a number of moles. A second calculation can be made assuming that one mole of glyphosate applied is converted into one mole of AMPA. The amount of AMPA equivalent from an application of GBH at a rate of 1.67 L·ha^-1^ with 540 g·L^-1^ active ingredient is of the order of 590 g·ha^-1^ AMPA equivalent. Considering the average AMPA content in MBS measured in this study (6.26 ug·g^-1^), we can see that an application of MBS at a rate of 4 t/ha^-1^ corresponds to 13 g·ha^-1^ of AMPA equivalent. Thus, one GBH application represents a potential maximum AMPA input to soil about 40 times greater than that that deriving from one 4 t·ha^-1^ MBS application (590 g·ha^-1^ / 13 g·ha^-1^).

To further validate our results on the use of MBS in agriculture, future studies should assess the longer-term impacts of MBS on glyphosate and AMPA contents in soils at a larger number of sites. It would be relevant to make an inventory of glyphosate and AMPA contents in MBS from different wastewater treatment plants and see whether, at different spreading rates, MBS could contribute significantly to glyphosate and AMPA contribution in soils. It will also be essential to determine the impact of MBS fertilisation in agricultural fields on the resident soil bacterial communities as we do our companion article (Blakney *et al*., 2023). Applying MBS to agricultural fields raises important questions about how MBS may change bacterial community composition, and function, or contribute human pathogenic bacteria to the food supply chain. We know that MBS can contain low contents of various potential human pathogens (Ryan *et al*., 2009; Walterson *et al*., 2015; Depoorter *et al*., 2016; Scott *et al*., 2022). With a pressing need for this data, our companion article, Blakney *et al*., (2023) investigated the impact of MBS use on bacterial communities in agricultural soils.

### 4.2 Risks of agricultural soil contamination by trace metals

Another objective of our study was to assess whether the application of MBS leads to an accumulation of trace metals in the soil. Trace metals have long been a source of concern for many agricultural stakeholders (Perron & Hébert, 2007). High contents of these elements can be toxic for crops (Hébert, 2015). In order to ensure product quality, the Québec regulation limits trace elements contents in MBS. Our results did not show any significant differences in soil trace elements contents between control plots and those treated with MBS (Figure S5). On the other hand, several studies from different countries, have reported higher trace metals contents due to MBS application (Marcela *et al*., 2020; Perron & Hébert, 2008; Pepper *et al*., 2008). However, the contents of these elements were systematically below the limits defined by each country.

### 4.3 MBS are a substitute to mineral fertilisers with agronomic, economic and environmental benefits

MBS are rich in N and P, and their use can provide competitive yields for both corn and soybean. Indeed, several studies have shown that fertilising crops with MBS provides yields as good as with mineral fertilisers (Vasseur *et al*., 1999; Warman & Termeer, 2005; Hébert, 2015). Our results point to the same trend, with equivalent yields between plots treated with MBS and plots receiving equivalent fertilization via mineral fertilisers only (Table 3). MBS application to agriculture soils represents a significant potential for providing nutrients and reducing mineral fertilisers, and thereby limit GHG emissions associated with MBS incineration or landfilling (Hébert, 2015). MBS applications to soils promote the degradation of OM under aerobic conditions, which are far less conducive to the formation of powerful GHG such as CH4 (Vigneux *et al*., 2016). Furthermore, although mineral fertilisers are an essential input for agricultural crops in Canada, the application of N fertilisers produces N2O, a GHG with a global warming potential 265 times greater than CO2 over a 100-year period (IPCC, 2022). The production and transport of fertilisers, particularly N fertilisers, also generates considerable amounts of GHG (Chai *et al*., 2019; Chataut *et al*., 2023). Growers also benefit economically from using MBS as fertilisers, as they can access them at low cost, and reducing the need to purchase expensive mineral fertilisers.

## 5 Conclusion

The ubiquity of glyphosate and AMPA in the environment is at the heart of hot debates around the world. Meanwhile, the agriculture sector needs to be increasingly resilient in the face of the climate crisis. Farmers are being asked to make greater use of organic fertilisers and residual fertilising materials to reduce their dependence on mineral fertilisers, which are monetarily and energetically costly and contribute to GHG emissions. Therefore, municipal biosolids could be an alternative to conventional mineral fertilisers.

Our study enabled a better understanding of the environmental and economic values associated with using MBS as fertilisers. Glyphosate and AMPA contents in soils of experimental field plots treated with MBS were measured and compared to control plots fertilised with mineral fertiliser. Although glyphosate and AMPA were measured in significant quantities in MBS, there were no differences in glyphosate and AMPA soil contents between treatments. In addition, corn and soybean yields were as high in MBS treated plots as in control plots. As such, from an environmental point of view, MBS is a convincing alternative to mineral fertilisers. Their use reduces GHG emissions linked to incineration and landfilling of wastewater treatment by-products, in addition to limiting GHG emissions associated with mineral fertiliser production. To our knowledge, this is one of the first reports presenting the contents of these two compounds in soil following MBS application. In the future, it would be interesting to assess glyphosate and AMPA contents in MBS from different wastewater treatment plants, to expand to other locations, and over longer time-scales.

## Supporting information

Charbonneau et al., 2023 BioRxiv Supplemental Materials

## Acknowledgements

This work was supported by the Ministry of Agriculture, Fisheries and Food of Quebec to M.L. (Grant Numbers: IA119565), which are gratefully acknowledged.

## Author Contributions

All authors contributed to the study conception and design. Funding acquisition as well as the provision of resources were provided by M.L. (Marc Lucotte). A.C. was supervised by M.L. (Marc Lucotte) and M.M. (Matthieu Moingt). Material preparation, data collection and analyses were performed by A.C., M.M. Soil sampling was realized by A.C. and M.M. The first draft of the paper was written by A.C. and all authors commented on previous versions of the paper. All authors have read and agreed to the published version of the paper.

## Informed Consent Statement

Informed consent was obtained from all subjects involved in the study.

## Data Availability Statement

The data presented in this study are available within the article or on request from Marc Lucotte.

