## Supplementary material for "Fertilisation of agricultural soils with municipal biosolids: Part 1- Glyphosate and aminomethylphosphonic acid inputs to Québec field crop soils": Charbonneau et al., 2023 BioRxiv Supplemental Materials

Table S1. 2021 agricultural management calendar.

| Site | Crop | Cultivar | Pre-seeding herbicide | | | Post-seeding herbicide #1 | | | Post-seeding herbicide #2 | | | Fertilizer #1 | | | Fertilizer #2 | | | Biosolids | |
|---|---|---|---|---|---|---|---|---|---|---|---|---|---|---|---|---|---|---|---|
| | | | **Date** | **Product** | **Dose (L/ha)** | **Date** | **Product** | **Dose (L/ha)** | **Date** | **Product** | **Dose (L/ha)** | **Date** | **Formulation** | **Dose (kg/ha)** | **Date** | **Formulation** | **Dose (kg/ha)** | **Date** | **Dose (ton/ha)** |
| CÉROM | Corn | MZ3690DBR | May 4 | Weather Max | 3.3 | June 9 | WeatherMax | 3 | n.a. | | | May 11 | 27-0-0 | 185 | June 7 | 46-0-0 | 125 | May 11 | 1 |
| | | | | | | | | | | | | | 0-46-0 | 141 | | | | | |
| | Soy | Maris R2X | April 29 | Weather Max | 3.3 | | WeatherMax | 2.5 | | | | | 0-46-0 | 130 | n.a. | | | | |
| | | | | | | | Extendimax | 0.825 | | | | | | | | | | | |
| La Présentation | Corn | MZ3818 | April 14 | Weather Max | 2.5 | May 12 | WeatherMax | 1.2 | June 10 | Weather Max | 2.5 | May 9 | 7-25-3 | 60* | June 2 | 46-0-0 | 55 | May 4 | 2.02 |
| | | | | | | | Integrity | 0.9 | | | | | 32-0-0 | 125* | | | | | |
| | | | | | | | | | | | | | 21-0-0 | 45* | | | | | |
| | Soy | Cobra R2X | n.a. | | | May 15 | WeatherMax | 3.3 | June 29 | Weather Max | 2.2 | May 15 | 7-25-3 | 30* | n.a. | | | | 3.78 |
| | | | | | | | Frontier | 0.8 | | | | | | | | | | | |
| St-Césaire | Corn | Pride 6012G2 | April 29 | Crucial 540 | 1 | May 24 | StoneWall 540 | 2.5 | n.a. | | | May 11 | 25-0-0 | 260 | June 28 | 32-0-0 | 150* | May 8 | 2.1 |
| | | | | | | | Destra IS | 275 g/ha | | | | | 6-24-6 | 50* | | | | | |
| | | | | | | | Lontrel XC | 0.167 | | | | | | | | | | | |
| | Soy | Ezra | n.a. | | | May 15 | Fortran 540 | 2.5 | June 21 | Arrow-X-Act | 1.125 | May 14 | 6-24-6 | 50* | n.a. | | | | |
| | | | | | | | Diligent | 176 g/ha | | | | | | | | | | | |
| | | | | | | | Pursuit | 0.31 | | | | | | | | | | | |
| St-Robert | Corn | DK-4617 | n.a. | | | June 10 | CreditXtreme | 3 | n.a. | | | May 11 | 19-19-0 | 188* | June 23 | 32-0-0 | 233* | May 13 | 6.3 |
| | | | | | | | | | | | | | 20--8-24 | 248 | | | | | |
| | Soy | Pionner P09A53X | | | | | | | | | | n.a. | | | n.a. | | | | |

*, L/ha

Table S2.  2022 agricultural management calendar.

| Site | Crop | Cultivar | Pre-seeding herbicide | Post-seeding herbicide #1 | | | Post-seeding herbicide #2 | | | Fertilizer #1 | | | Fertilizer #2 | | | Biosolids | |
|---|---|---|---|---|---|---|---|---|---|---|---|---|---|---|---|---|---|
| | | | | Date | Product | Dose (L/ha) | Date | Product | Dose (L/ha) | Date | Formulation | Dose (kg/ha) | Date | Formulation | Dose (kg/ha) | Date | Dose (ton/ha) |
| CÉROM | Corn | MZ3505DBR | n.a. | June 14 | Factor 540 | 3 | n.a. | | | May 12 | 18-18-04 | 270 | June 3 | 46-0-0 | 260 | May 11 | 6.6 |
| | Soy | P12T94E | | | Roundup Transorb HC | 2.3 | | | | June 4 | 0-46-0 | 20 | n.a. | | | | 3.5 |
| La Présentation | Corn | MZ3117 | n.a. | May 7 | WeatherMax | 2.4 | June 20 | WeatherMax | 2.7 | May 7 | 7-25-3 | 60* | June 3 | 46-0-0 | 55 | May 6 | 2.04 |
| | | | | May 12 | Integrity | 0.9 | | | | | 32-0-0 | 125* | | | | | |
| | | | | | | | | | | | 21-0-0 | 45* | | | | | |
| | Soy | Kytes | | May 11 | WeatherMax | 3.3 | June 20 | Enlist Duo | 4.34 | n.a. | | | n.a. | | | | |
| | | | | | Frontier | 0.8 | | | | | | | | | | | |
| St-Césaire | Corn | 3490VT2P | n.a. | May 25 | Engenia | 0.74 | n.a. | | | May 12 | 25-0-0 | 190 | June 21 | 32-0-0 | 204* | May 12 | 3.3 |
| | | | | | StoneWall 540 | 2.5 | | | | | 6-24-6 | 50* | | | | | |
| | | | | | Destra IS | 275 g/ha | | | | | | | | | | | |
| | Soy | Hana | | May 18 | StoneWall 540 | 2.5 | | | | May 14 | 6-24-6 | 50* | n.a. | | | | 3.5 |
| | | | | | Boundary | 2.5 | | | | | | | | | | | |
| | | | | | Broadstrike | 90 g/ha | | | | | | | | | | | |
| St-Robert | Corn | PC3575VT2 | n.a. | June 20 | CreditXtreme | 3 | n.a. | | | May 11 | 19-19-0 | 188* | June 23 | 32-0-0 | 233* | May 25 | 5.8 |
| | | | | | | | | | | | 20--8-24 | 248 | | | | | |
| | Soy | Pionner P09A53X | | | | | | | | n.a. | | | n.a. | | | | |

*, L/ha

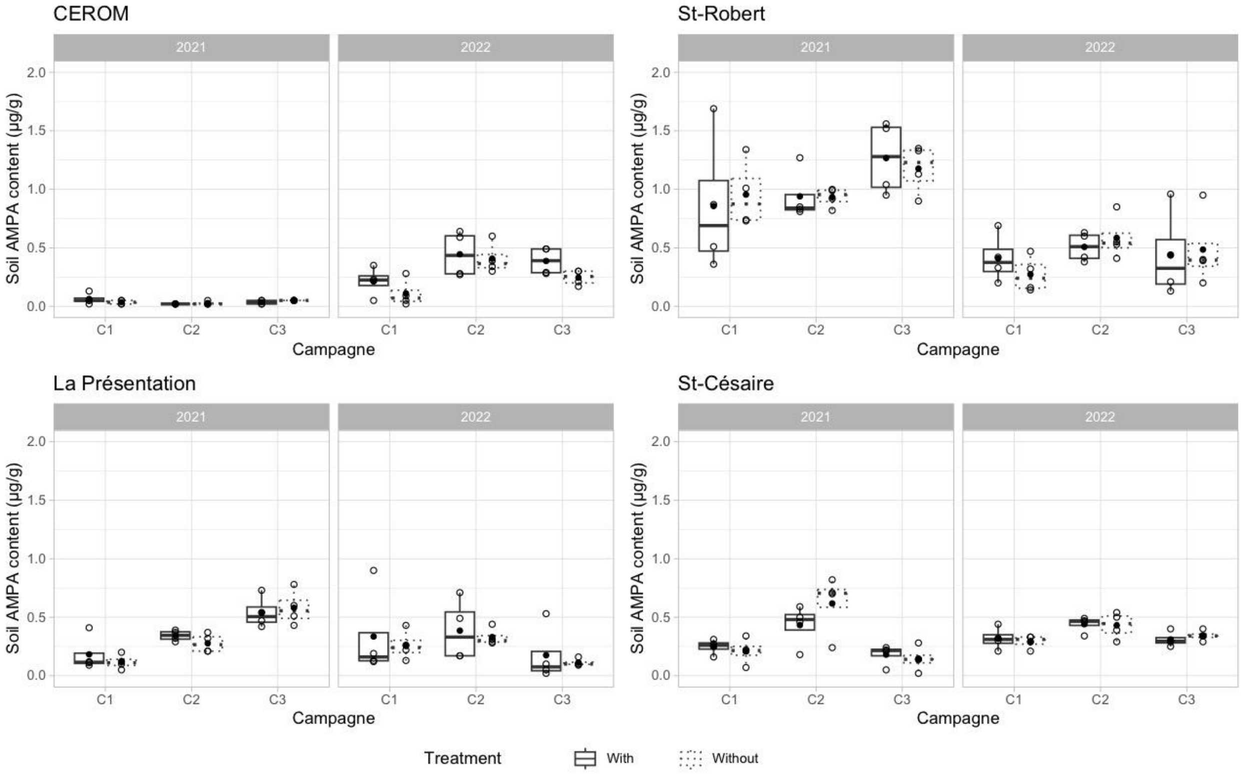


Figure S1. Soil AMPA content (µg/g) in corn fields at CEROM, at St-Robert, at La Présentation and at St-Césaire measured during each sampling campaign in 2021 and 2022 at CEROM, St-Robert, La Présentation and St-Césaire.

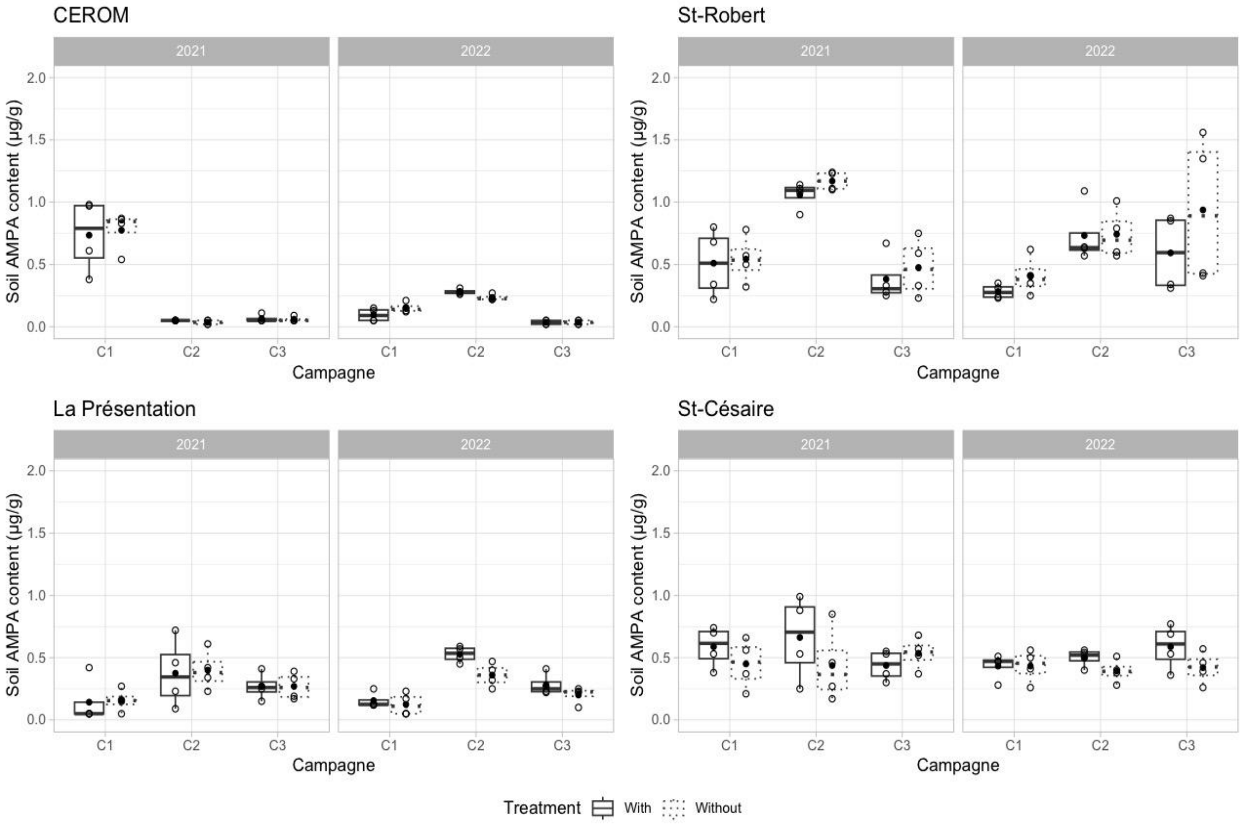


Figure S2. Soil AMPA content (µg/g) in soybean fields measured during each sampling campaign in 2021 and 2022 at CEROM, St-Robert, La Présentation and St-Césaire.

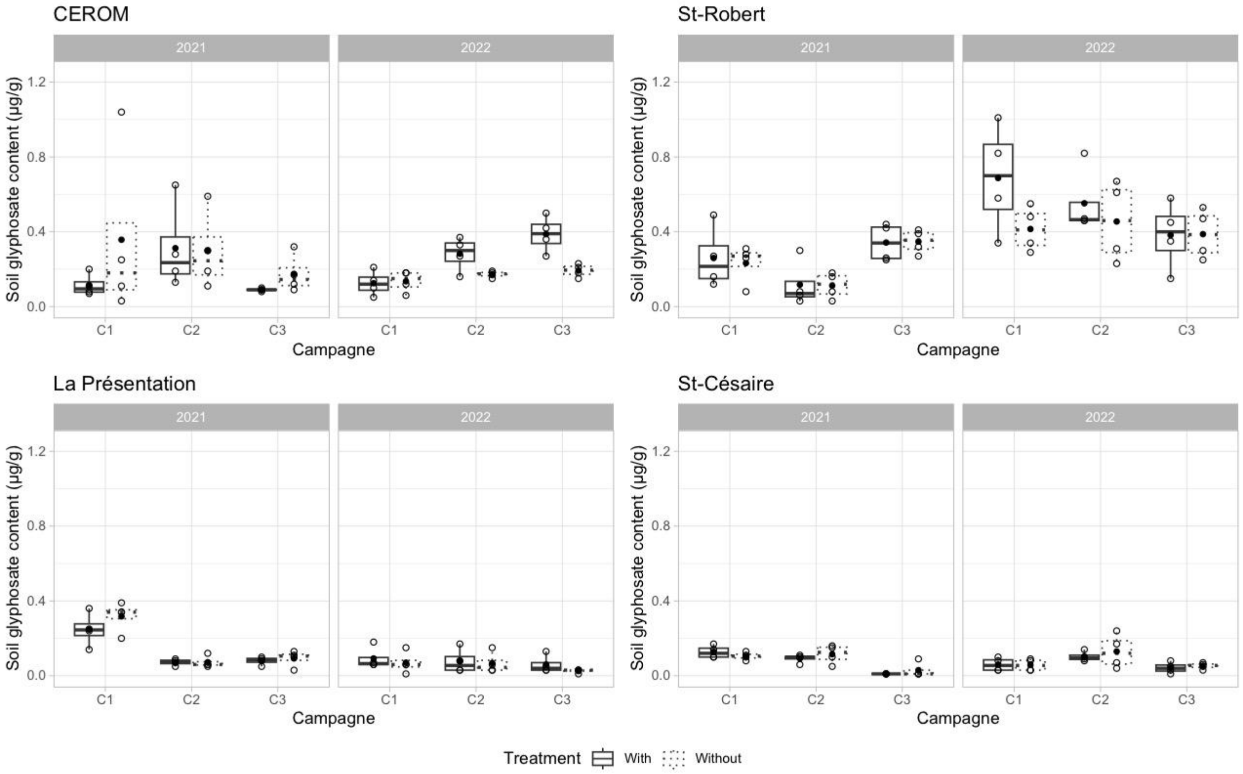


Figure S3. Soil glyphosate content (µg/g) in corn fields measured during each sampling campaign in 2021 and 2022 at CEROM, St-Robert, La Présentation and St-Césaire.

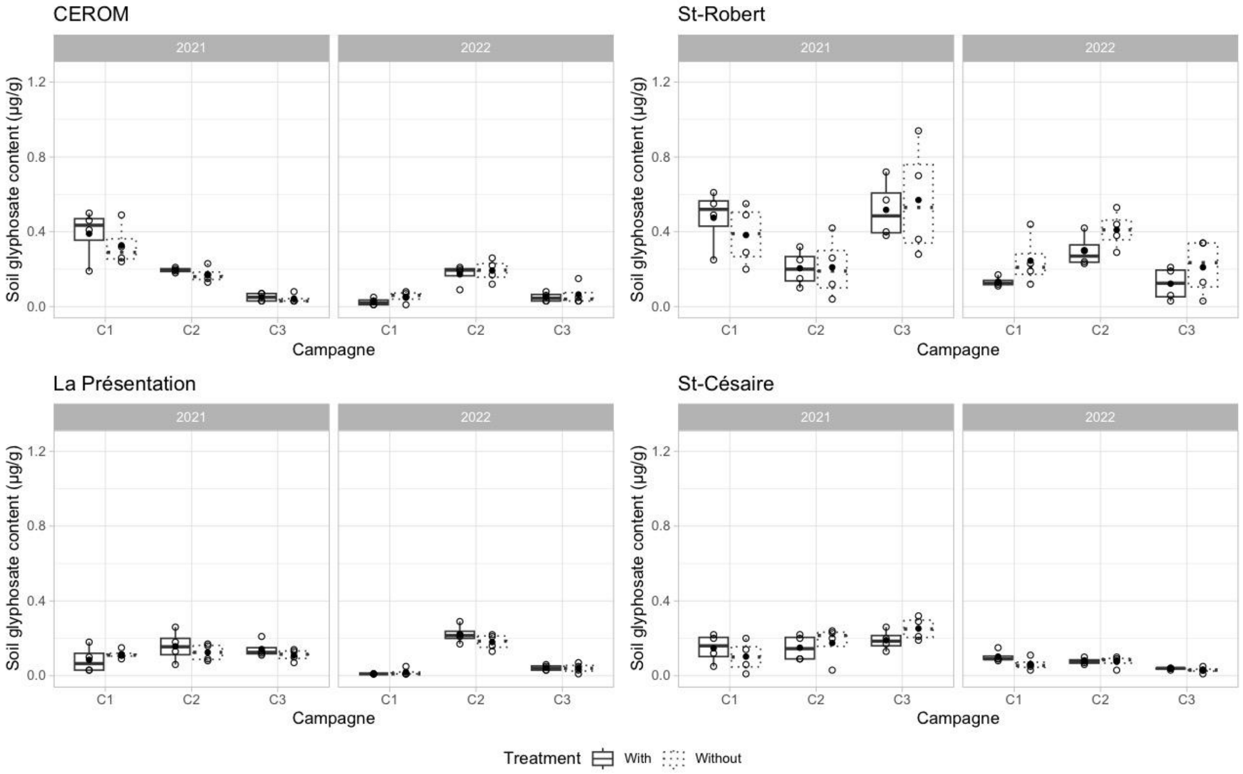


Figure S4. Soil glyphosate content (µg/g) in soybean fields measured during each sampling campaign in 2021 and 2022 at CEROM, St-Robert, La Présentation and St-Césaire.

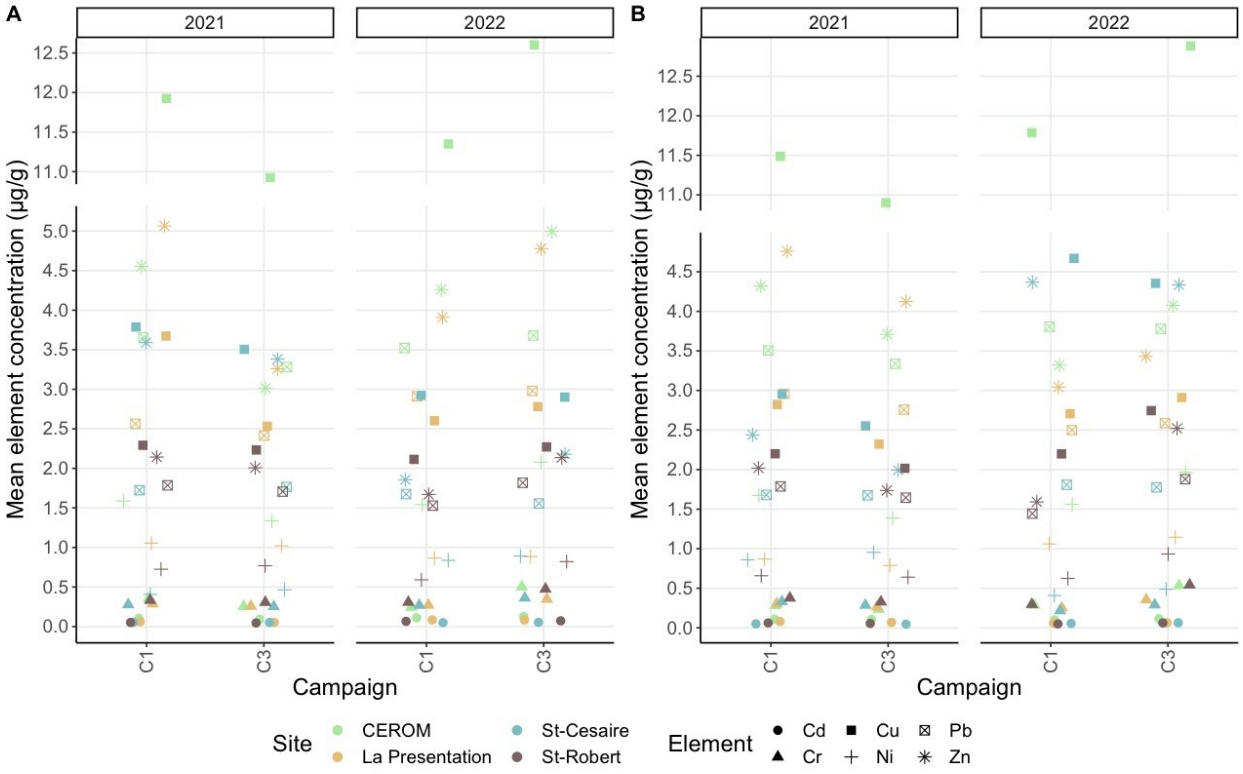


Figure S5. Soil element content (µg/g) in (A) corn and (B) soybean fields.

Table S1. 2021 agricultural management calendar.

| Site | Crop | Cultivar | Pre-seeding herbicide | | | Post-seeding herbicide #1 | | | Post-seeding herbicide #2 | | | Fertilizer #1 | | | Fertilizer #2 | | | Biosolids | |
|---|---|---|---|---|---|---|---|---|---|---|---|---|---|---|---|---|---|---|---|
| | | | Date | Product | Dose (L/ha) | Date | Product | Dose (L/ha) | Date | Product | Dose (L/ha) | Date | Formulation | Dose (kg/ha) | Date | Formulation | Dose (kg/ha) | Date | Dose (ton/ha) |
| CÉROM | Corn | MZ3690DBR | May 4 | Weather Max | 3.3 | June 9 | WeatherMax | 3 | n.a. | | | May 11 | 27-0-0 | 185 | June 7 | 46-0-0 | 125 | May 11 | 1 |
| | | | | | | | | | | | | | 0-46-0 | 141 | | | | | |
| | Soy | Maris R2X | April 29 | Weather Max | 3.3 | | WeatherMax | 2.5 | | | | | 0-46-0 | 130 | n.a. | | | | |
| | | | | | | | Extendimax | 0.825 | | | | | | | | | | | |
| La Présentation | Corn | MZ3818 | April 14 | Weather Max | 2.5 | May 12 | WeatherMax | 1.2 | June 10 | Weather Max | 2.5 | May 9 | 7-25-3 | 60* | June 2 | 46-0-0 | 55 | May 4 | 2.02 |
| | | | | | | | Integrity | 0.9 | | | | | 32-0-0 | 125* | | | | | |
| | | | | | | | | | | | | | 21-0-0 | 45* | | | | | |
| | Soy | Cobra R2X | n.a. | | | May 15 | WeatherMax | 3.3 | June 29 | Weather Max | 2.2 | May 15 | 7-25-3 | 30* | n.a. | | | | 3.78 |
| | | | | | | | Frontier | 0.8 | | | | | | | | | | | |
| St-Césaire | Corn | Pride 6012G2 | April 29 | Crucial 540 | 1 | May 24 | StoneWall 540 | 2.5 | n.a. | | | May 11 | 25-0-0 | 260 | June 28 | 32-0-0 | 150* | May 8 | 2.1 |
| | | | | | | | Destra IS | 275 g/ha | | | | | 6-24-6 | 50* | | | | | |
| | | | | | | | Lontrel XC | 167 g/ha | | | | | | | | | | | |
| | Soy | Ezra | n.a. | | | May 15 | Fortran 540 | 2.5 | June 21 | Arrow-X-Act | 1.125 | May 14 | 6-24-6 | 50* | n.a. | | | | |
| | | | | | | | Diligent | 0.176 | | | | | | | | | | | |
| | | | | | | | Pursuit | 0.31 | | | | | | | | | | | |
| St-Robert | Corn | DK-4617 | n.a. | | | June 10 | CreditXtreme | 3 | n.a. | | | May 11 | 19-19-0 | 188* | June 23 | 32-0-0 | 233* | May 13 | 6.3 |
| | | | | | | | | | | | | | 20--8-24 | 248 | | | | | |
| | Soy | Pionner P09A53X | | | | | | | | | | n.a. | | | n.a. | | | | |

*, L/ha
